# Inhibitory effect of eslicarbazepine acetate and S-licarbazepine on Na_v_1.5 channels

**DOI:** 10.1101/2020.04.24.059188

**Authors:** Theresa K. Leslie, Lotte Brückner, Sangeeta Chawla, William J. Brackenbury

**Author notes:** Correspondence: Dr William J. Brackenbury, Department of Biology and York Biomedical Research Institute, University of York, Wentworth Way, Heslington, York YO10 5DD, UK.

## Abstract

Eslicarbazepine acetate (ESL) is a dibenzazepine anticonvulsant approved as adjunctive treatment for partial-onset epileptic seizures. Following first pass hydrolysis of ESL, S-licarbazepine (S-Lic) represents around 95 % of circulating active metabolites. S-Lic is the main enantiomer responsible for anticonvulsant activity and this is proposed to be through the blockade of voltage-gated Na^+^ channels (VGSCs). ESL and S-Lic both have a voltage-dependent inhibitory effect on the Na^+^ current in N1E-115 neuroblastoma cells expressing neuronal VGSC subtypes including Na_v_1.1, Na_v_1.2, Na_v_1.3, Na_v_1.6 and Na_v_1.7. ESL has not been associated with cardiotoxicity in healthy volunteers, although a prolongation of the electrocardiographic PR interval has been observed, suggesting that ESL may also inhibit cardiac Na_v_1.5 isoform. However, this has not previously been studied. Here, we investigated the electrophysiological effects of ESL and S-Lic on Na_v_1.5 using whole-cell patch clamp recording. We interrogated two model systems: (1) MDA-MB-231 metastatic breast carcinoma cells, which endogenously express the ‘neonatal’ Na_v_1.5 splice variant, and (2) HEK-293 cells stably over-expressing the ‘adult’ Na_v_1.5 splice variant. We show that both ESL and S-Lic inhibit transient and persistent Na^+^ current, hyperpolarise the voltage-dependence of fast inactivation, and slow the recovery from channel inactivation. These findings highlight, for the first time, the potent inhibitory effects of ESL and S-Lic on the Na_v_1.5 isoform, suggesting a possible explanation for the prolonged PR interval observed in patients on ESL treatment. Given that numerous cancer cells have also been shown to express Na_v_1.5, and that VGSCs potentiate invasion and metastasis, this study also paves the way for future investigations into ESL and S-Lic as potential invasion inhibitors.

## 1 Introduction

Eslicarbazepine acetate (ESL) is a member of the dibenzazepine anticonvulsant family of compounds which also includes oxcarbazepine and carbamazepine (1). ESL has been approved by the European Medicines Agency and the United States Federal Drug Administration as an adjunctive treatment for partial-onset epileptic seizures (2). ESL is administered orally and rapidly undergoes first pass hydrolysis to two stereoisomeric metabolites, R-licarbazepine and S-licarbazepine (S-Lic; also known as eslicarbazepine; Figure 1A, B) (3-5). S-Lic represents around 95 % of circulating active metabolites following first pass hydrolysis of ESL and is the enantiomer responsible for anticonvulsant activity (6, 7). S-Lic also has improved blood brain barrier penetration compared to R-licarbazepine (8). Although S-Lic has been shown to inhibit T type Ca^2+^ channels (9), its main activity is likely through blockade of voltage-gated Na^+^ channels (VGSCs) (10). ESL offers several clinical advantages over other older VGSC-inhibiting antiepileptic drugs, e.g. carbamazepine, phenytoin; it has a favourable safety profile (10, 11), reduced induction of hepatic cytochrome P450 enzymes (12), low potential for drug-drug interactions (13, 14), and takes less time to reach a steady state plasma concentration (15).

**Figure 1.**
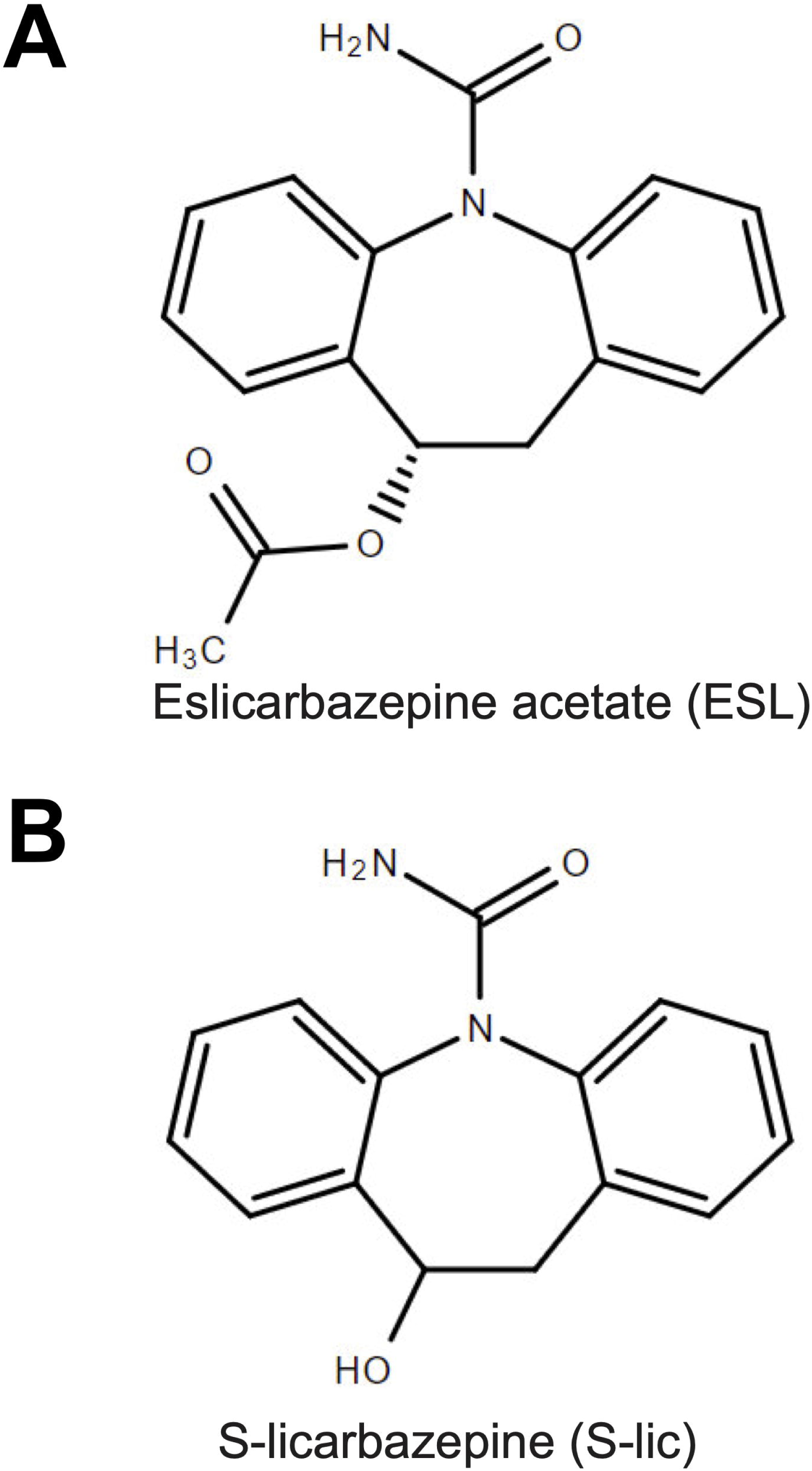
Chemical structures of eslicarbazepine acetate and S-licarbazepine. (A) eslicarbazepine acetate; (9S)-2-carbamoyl-2-azatricyclo[9.4.0.0^3,^□]pentadeca-1(15),3,5,7,11,13-hexaen-9-yl acetate. (B) S-licarbazepine; (10R)-10-hydroxy-2-azatricyclo[9.4.0.0^3,8^]pentadeca-1(11),3,5,7,12,14-hexaene-2-carboxamide. Structures were drawn using Chemspider software.

VGSCs are composed of a pore-forming α subunit in association with one or more auxiliary β subunits, the latter modulating channel gating and kinetics in addition to functioning as cell adhesion molecules (16). There are nine α subunits (Na_v_1.1-Na_v_1.9), and four β subunits (β1-4) (17, 18). In postnatal and adult CNS neurons, the predominant α subunits are the tetrodotoxin-sensitive Na_v_1.1, Na_v_1.2 and Na_v_1.6 isoforms (19) and it is therefore on these that the VGSC-inhibiting activity of ESL and S-Lic has been described. In the murine neuroblastoma N1E-115 cell line, which expresses Na_v_1.1, Na_v_1.2, Na_v_1.3, Na_v_1.6 and Na_v_1.7, ESL and S-Lic both have a voltage-dependent inhibitory effect on the Na^+^ current (10, 20). In this cell model, S-Lic has no effect on the voltage-dependence of fast inactivation, but significantly hyperpolarises the voltage-dependence of slow inactivation (10). S-Lic also has a lower affinity for VGSCs in the resting state than carbamazepine or oxcarbazepine, thus potentially improving its therapeutic window over first- and second-generation dibenzazepine compounds (10). In acutely isolated murine hippocampal CA1 neurons, which express Na_v_1.1, Na_v_1.2 and Na_v_1.6 (21-23), S-Lic significantly reduces the persistent Na^+^ current, a very slow-inactivating component ∼1 % the size of the peak transient Na^+^ current (24, 25). Moreover, in contrast to carbamazepine, this effect is maintained in the absence of β1 (24, 26).

In healthy volunteers, ESL has not been associated with cardiotoxicity and the QT interval remains unchanged on treatment (27). However, a prolongation of the PR interval has been observed (27), suggesting that caution should be exercised in patients with cardiac conduction abnormalities (13). Prolongation of the PR interval suggests that ESL may also inhibit the cardiac Na_v_1.5 isoform, although this has not previously been studied. Na_v_1.5 is not only responsible for the initial depolarisation of the cardiac action potential (28), but is also expressed in breast and colon carcinoma cells, where the persistent Na^+^ current promotes invasion and metastasis (29-32). Inhibition of Na_v_1.5 with phenytoin or ranolazine decreases tumour growth, invasion and metastasis (33-35). Thus, it is of interest to specifically understand the effect of ESL on the Na_v_1.5 isoform.

In the present study we investigated the electrophysiological effects of ESL and S-Lic on Na_v_1.5 [1] endogenously expressed in the MDA-MB-231 metastatic breast carcinoma cell line, and [2] stably over-expressed in HEK-293 cells. We show that both ESL and S-Lic inhibit transient and persistent Na^+^ current, hyperpolarise the voltage-dependence of fast inactivation, and slow the recovery from channel inactivation. These findings highlight, for the first time, the potent inhibitory effects of ESL and S-Lic on the Na_v_1.5 isoform.

## 2 Materials and methods

### 2.1 Pharmacology

ESL (Tokyo Chemical Industry UK Ltd) was dissolved in DMSO to make a stock concentration of 67 mM. S-Lic (Tocris) was dissolved in DMSO to make a stock concentration of 300 mM. Both drugs were diluted to working concentrations of 100-300 µM in extracellular recording solution. The concentration of DMSO in the recording solution was 0.45 % for ESL and 0.1 % for S-Lic. Equal concentrations of DMSO were used in the control solutions. DMSO (0.45 %) had no effect on the Na^+^ current (Supplementary Figure 1).

### 2.2 Cell culture

MDA-MB-231 cells and HEK-293 cells stably expressing Na_v_1.5 (a gift from L. Isom, University of Michigan) were grown in Dulbecco’s modified eagle medium supplemented with 5 % FBS and 4 mM L-glutamine (36). Molecular identity of the MDA-MB-231 cells was confirmed by short tandem repeat analysis (37). Cells were confirmed as mycoplasma-free using the DAPI method (38). Cells were seeded onto glass coverslips 48 h before electrophysiological recording.

### 2.3 Electrophysiology

Plasma membrane Na^+^ currents were recorded using the whole-cell patch clamp technique, using methods described previously (32, 35). Patch pipettes made of borosilicate glass were pulled using a P-97 pipette puller (Sutter Instrument) and fire-polished to a resistance of 3-5 MΩ when filled with intracellular recording solution. The extracellular recording solution for MDA-MB-231 cells contained (in mM): 144 NaCl, 5.4 KCl, 1 MgCl_2_, 2.5 CaCl_2_, 5.6 D-glucose and 5 HEPES (adjusted to pH 7.2 with NaOH). For the extracellular recording solution for HEK-293 cells expressing Na_v_1.5, the extracellular [Na^+^] was reduced to account for the much larger Na^+^ currents and contained (in mM): 60 NaCl, 84 Choline Cl, 5.4 KCl, 1 MgCl_2_, 2.5 CaCl_2_, 5.6 D-glucose and 5 HEPES (adjusted to pH 7.2 with NaOH). The intracellular recording solution contained (in mM): 5 NaCl, 145 CsCl, 2 MgCl_2_, 1 CaCl_2_, 10 HEPES, 11 EGTA, (adjusted to pH 7.4 with CsOH) (39). Voltage clamp recordings were made at room temperature using a Multiclamp 700B or Axopatch 200B amplifier (Molecular Devices) compensating for series resistance by 40–60%. Currents were digitized using a Digidata interface (Molecular Devices), low pass filtered at 10 kHz, sampled at 50 kHz and analysed using pCLAMP 10.7 software (Molecular Devices). Leak current was subtracted using a P/6 protocol (40). Extracellular recording solution ± drugs was applied to the recording bath at a rate of ∼1.5 ml/min using a ValveLink 4-channel gravity perfusion controller (AutoMate Scientific). Each new solution was allowed to equilibrate in the bath for ∼4 min following switching prior to recording at steady state.

### 2.4 Voltage clamp protocols

Cells were clamped at a holding potential of −120 mV or −80 mV for ≥ 250 ms, dependent on experiment (detailed in the Figure legends). Five main voltage clamp protocols were used, as follows:

1. To assess the effect of drug perfusion and wash-out on peak current in real time, a simple one-step protocol was used where cells were held at −120 mV or −80 mV for 250 ms and then depolarised to −10 mV for 50 ms.
2. To assess the voltage-dependence of activation, cells were held at −120 mV for 250 ms and then depolarised to test potentials in 10 mV steps between −120 mV and +30 mV for 50 ms. The voltage of activation was taken as the most negative voltage which induced a visible transient inward current.
3. To assess the voltage-dependence of steady-state inactivation, cells were held at −120 mV for 250 ms followed by prepulses for 250 ms in 10 mV steps between −120 mV and +30 mV and a test pulse to −10 mV for 50 ms.
4. To assess recovery from fast inactivation, cells were held at −120 mV for 250 ms, and then depolarised twice to 0 mV for 25 ms, returning to −120 mV for the following intervals between depolarisations (in ms): 1, 2, 3, 5, 7, 10, 15, 20, 30, 40, 50, 70, 100, 150, 200, 250, 350, 500. In each case, the second current was normalised to the initial current and plotted against the interval time.

### 2.5 Curve fitting and data analysis

To study the voltage-dependence of activation, current-voltage (I-V) relationships were converted to conductance using the following equation:

G = I / (V_m_ – V_rev_), where G is conductance, I is current, V_m_ is the membrane voltage and V_rev_ is the reversal potential for Na^+^ derived from the Nernst equation. Given the different recording solutions used, V_rev_ for Na^+^ was +85 mV for MDA-MB-231 cells and +63 mV for HEK-Na_v_1.5 cells.

The voltage-dependence of conductance and availability were normalised and fitted to a Boltzmann equation:

G = G_max_ / (1 + exp ((V_1/2_ – V_m_) / k)), where G_max_ is the maximum conductance, V_1/2_ is the voltage at which the channels are half activated/inactivated, V_m_ is the membrane voltage and k is the slope factor.

Recovery from inactivation data (I_t_ / I_t=0_) were normalised, plotted against recovery time (Δt) and fitted to a single exponential function:

τ = A_1_ + A_2_ exp (-t / t_0_), where A_1_ and A_2_ are the coefficients of decay of the time constant (τ), t is time and t_0_ is a time constant describing the time dependence of τ.

The time course of inactivation was fitted to a double exponential function:

I = A_f_ exp (-t / τ_f_) + A_s_ exp (-t / τ_s_) + C, where A_f_ and A_s_ are maximal amplitudes of the slow and fast components of the current, τ_f_ and τ_s_ are the fast and slow decay time constants and C is the asymptote.

### 2.6 Statistical analysis

Data are presented as mean and SEM unless stated otherwise. Statistical analysis was performed on the raw (non-normalised) data using GraphPad Prism 8.4.0. Pairwise statistical significance was determined with Student’s paired *t*-tests. Multiple comparisons were made using ANOVA and Tukey post-hoc tests, unless stated otherwise. Results were considered significant at *P* < 0.05.

## 3 Results

### 3.1 Effect of eslicarbazepine acetate and S-licarbazepine on transient and persistent Na^+^ current

Several studies have clearly established the inhibition of neuronal VGSCs (Na_v_1.1, Na_v_1.2, Na_v_1.3, Na_v_1.6, Na_v_1.7 and Na_v_1.8) by ESL and its active metabolite S-Lic (10, 20, 24, 41). Given that ESL prolongs the PR interval (27), potentially via inhibiting the cardiac Na_v_1.5 isoform, together with the interest in inhibiting Na_v_1.5 in carcinoma cells to reduce invasion and metastasis (33, 34, 42-44), it is also relevant to evaluate the electrophysiological effects of ESL and S-Lic on this isoform. We therefore evaluated the effect of both compounds on Na_v_1.5 current properties using whole-cell patch clamp recording, employing a two-pronged approach: (1) recording Na_v_1.5 currents endogenously expressed in the MDA-MB-231 breast cancer cell line (29, 30, 45), and (2) recording from Na_v_1.5 stably over-expressed in HEK-293 cells (HEK-Na_v_1.5) (46).

Initially, we evaluated the effect of both compounds on the size of the peak Na^+^ current in MDA-MB-231 cells. Na^+^ currents were elicited by depolarising the membrane potential (V_m_) to −10 mV from a holding potential (V_h_) of −120 mV or −80 mV. Application of the prodrug ESL (300 μM) reversibly inhibited the transient Na^+^ current by 49.6 ± 3.2 % when the V_h_ was −120 mV (P < 0.001; n = 13; ANOVA + Tukey test; Figure 2A, D). When V_h_ was set to −80 mV, ESL (300 μM) reversibly inhibited the transient Na^+^ current by 79.5 ± 4.5 % (P < 0.001; n = 12; ANOVA + Tukey test; Figure 2C, E). We next assessed the effect of ESL in HEK-Na_v_1.5 cells. Application of ESL (300 μM) inhibited Na_v_1.5 current by 74.7 ± 4.3 % when V_h_ was −120 mV (P < 0.001; n = 12; Figure 2F, I) and by 90.5 ± 2.8 % when V_h_ was −80 mV (P < 0.001; n = 14; Figure 2H, J). However, the inhibition was only partially reversible (P < 0.001; n = 14; Figure 2F, H-J). Application of ESL at a lower concentration (100 µM) elicited a similar result (Supplementary Figure 2A-J & Supplementary Table 1). Together, these data suggest that ESL preferentially inhibited Na_v_1.5 in the open or inactivated state, since the current inhibition was greater at more depolarised V_h_.

**Figure 2.**
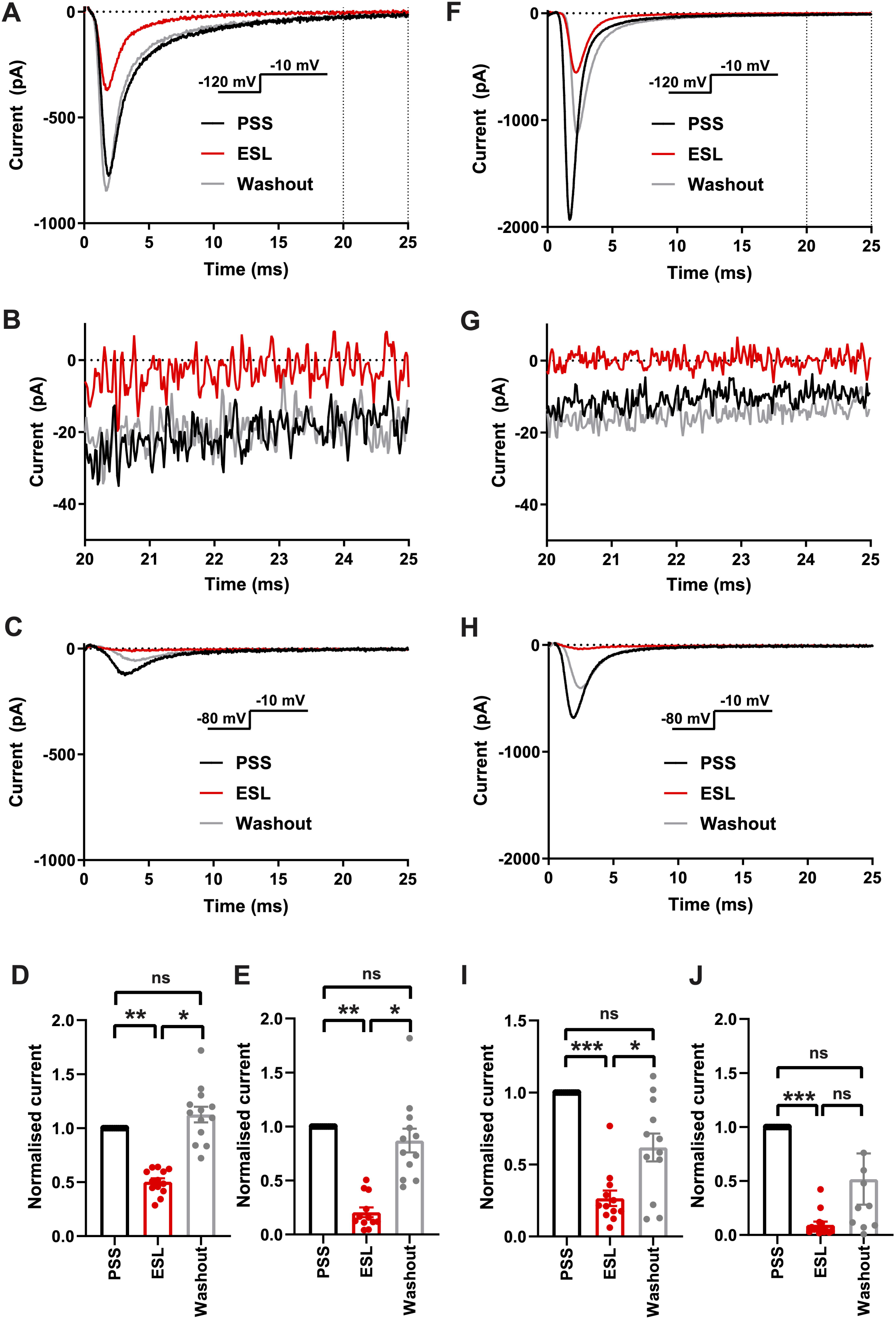
Effect of eslicarbazepine acetate on Na_v_1.5 currents. (A) Representative Na^+^ currents in an MDA-MB-231 cell elicited by a depolarisation from −120 mV to −10 mV in physiological saline solution (PSS; black), eslicarbazepine acetate (ESL; 300 μM; red) and after washout (grey). Dotted vertical lines define the time period magnified in (B). (B) Representative persistent Na^+^ currents in an MDA-MB-231 cell elicited by a depolarisation from −120 mV to −10 mV. (C) Representative Na^+^ currents in an MDA-MB-231 cell elicited by a depolarisation from −80 mV to −10 mV. (D) Normalised Na^+^ currents in MDA-MB-231 cells elicited by a depolarisation from −120 mV to −10 mV. (E) Normalised Na^+^ currents in MDA-MB-231 cells elicited by a depolarisation from −80 mV to −10 mV. (F) Representative Na^+^ currents in a HEK-Na_v_1.5 cell elicited by a depolarisation from −120 mV to −10 mV in PSS (black), ESL (300 μM; red) and after washout (grey). Dotted vertical lines define the time period magnified in (G). (G) Representative persistent Na^+^ currents in a HEK-Na_v_1.5 cell elicited by a depolarisation from −120 mV to −10 mV. (H) Representative Na^+^ currents in a HEK-Na_v_1.5 cell elicited by a depolarisation from −80 mV to −10 mV. (I) Normalised Na^+^ currents in HEK-Na_v_1.5 cells elicited by a depolarisation from −120 mV to −10 mV. (J) Normalised Na^+^ currents in HEK-Na_v_1.5 cells elicited by a depolarisation from −80 mV to −10 mV. Results are mean + SEM. *P ≤ 0.05; **P ≤ 0.01; ***P ≤ 0.001; one-way ANOVA with Tukey tests (n = 12-14). NS, not significant.

We next tested the effect of the active metabolite S-Lic. S-Lic (300 μM) inhibited the transient Na^+^ current in MDA-MB-231 cells by 44.4 ± 6.1 % when the V_h_ was −120 mV (P < 0.001; n = 9; ANOVA + Tukey test; Figure 3A, D). When V_h_ was set to −80 mV, S-Lic (300 µM) inhibited the transient Na^+^ current by 73.6 ± 4.1 % (P < 0.001; n = 10; ANOVA + Tukey test; Figure 3C, E). However, the inhibition caused by S-Lic (300 μM) was only partially reversible (P < 0.05; n = 10; ANOVA + Tukey test; Figure 3A, C-E). In HEK-Na_v_1.5 cells, S-Lic (300 μM) inhibited Na_v_1.5 current by 46.4 ± 3.9 % when V_h_ was −120 mV (P < 0.001; n = 13; ANOVA + Tukey test; Figure 3F, I) and by 74.0 ± 4.2 % when V_h_ was −80 mV (P < 0.001; n = 12; ANOVA + Tukey test; Figure 3H, J). Furthermore, the inhibition in HEK-Na_v_1.5 cells was not reversible over the duration of the experiment. Application of S-Lic at a lower concentration (100 µM) elicited a broadly similar result (Supplementary Figure 3A-J & Supplementary Table 1). Together, these data show that channel inhibition by S-Lic was also more effective at more depolarised V_h_. However, unlike ESL, channel blockade by S-Lic persisted after washout, suggesting higher target binding affinity for the active metabolite and/or greater trapping of the active metabolite in the cytoplasm.

**Figure 3.**
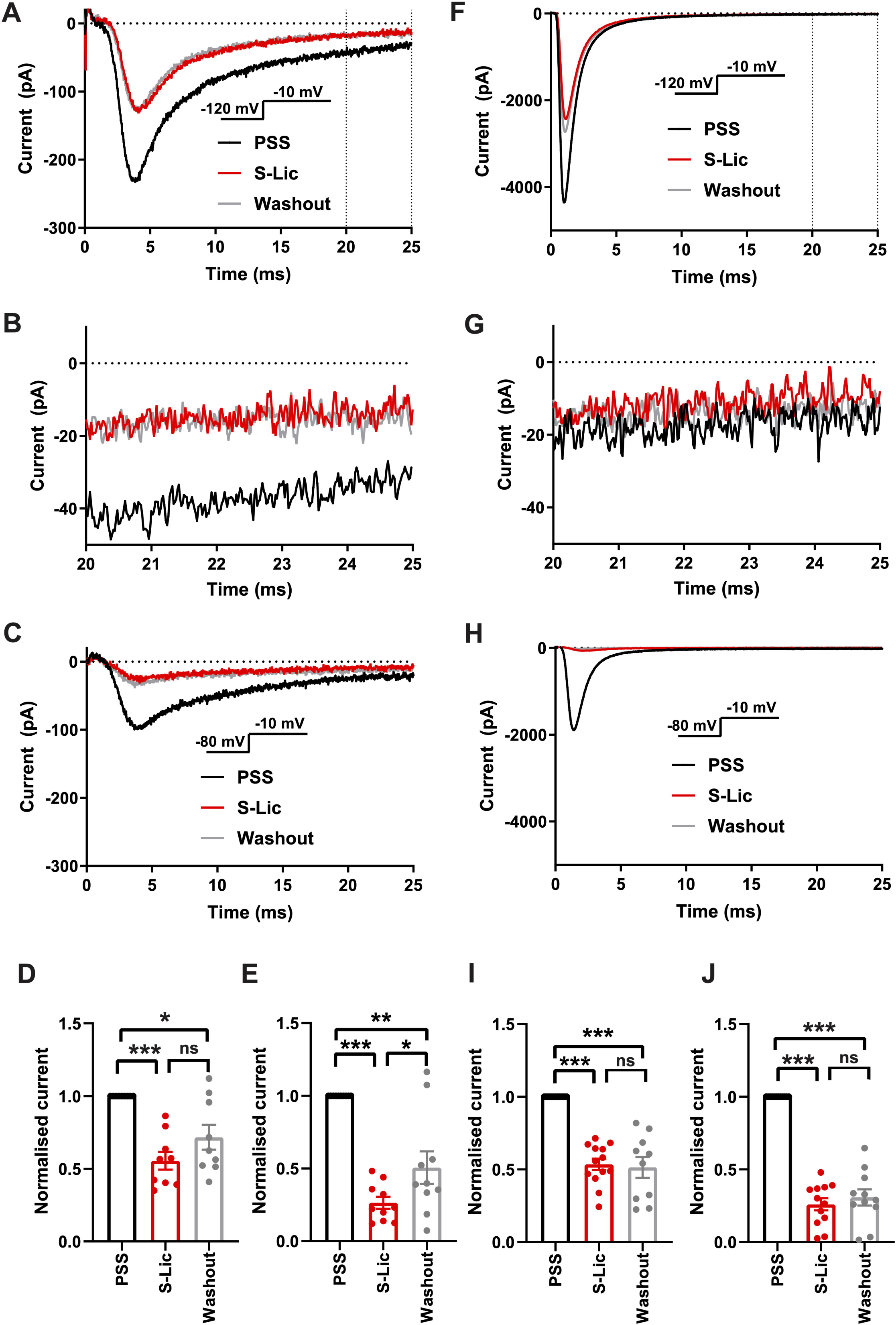
Effect of S-licarbazepine on Na_v_1.5 currents. (A) Representative Na^+^ currents in an MDA-MB-231 cell elicited by a depolarisation from −120 mV to −10 mV in physiological saline solution (PSS; black), S-licarbazepine (S-Lic; 300 μM; red) and after washout (grey). Dotted vertical lines define the time period magnified in (B). (B) Representative persistent Na^+^ currents in an MDA-MB-231 cell elicited by a depolarisation from −120 mV to −10 mV. (C) Representative Na^+^ currents in an MDA-MB-231 cell elicited by a depolarisation from −80 mV to −10 mV. (D) Normalised Na^+^ currents in MDA-MB-231 cells elicited by a depolarisation from −120 mV to −10 mV. (E) Normalised Na^+^ currents in MDA-MB-231 cells elicited by a depolarisation from −80 mV to −10 mV. (F) Representative Na^+^ currents in a HEK-Na_v_1.5 cell elicited by a depolarisation from −120 mV to - 10 mV in PSS (black), S-Lic (300 μM; red) and after washout (grey). Dotted vertical lines define the time period magnified in (G). (G) Representative persistent Na^+^ currents in a HEK-Na_v_1.5 cell elicited by a depolarisation from −120 mV to −10 mV. (H) Representative Na^+^ currents in a HEK-Na_v_1.5 cell elicited by a depolarisation from −80 mV to −10 mV. (I) Normalised Na^+^ currents in HEK-Na_v_1.5 cells elicited by a depolarisation from −120 mV to −10 mV. (J) Normalised Na^+^ currents in HEK-Na_v_1.5 cells elicited by a depolarisation from −80 mV to −10 mV. Results are mean + SEM. *P ≤ 0.05; ***P ≤ 0.001; one-way ANOVA with Tukey tests (n = 9-13). NS, not significant.

We also assessed the effect of both compounds on the persistent Na^+^ current measured 20-25 ms after depolarisation to −10 mV from −120 mV. In MDA-MB-231 cells, ESL (300 μM) inhibited the persistent Na^+^ current by 77 ± 34 % although the reduction was not statistically significant (P = 0.13; n = 12; paired t test; Figure 2B, Table 1). In HEK-Na_v_1.5 cells, ESL (300 μM) inhibited persistent current by 76 ± 10 % (P < 0.01; n = 12; paired t test; Figure 2G, Table 1). S-Lic (300 μM) inhibited the persistent Na^+^ current in MDA-MB-231 cells by 66 ± 16 % (P < 0.05; n = 9; paired t test; Figure 3B, Table 2). In HEK-Na_v_1.5 cells, S-Lic (300 μM) inhibited persistent current by 35 ± 16 % (P < 0.05; n = 11; Figure 3G, Table 2). Application of both compounds at a lower concentration (100 µM) elicited a similar result (Supplementary Table 1). In summary, both ESL and S-Lic also inhibited the persistent Na^+^ current.

**Table 1.** Effect of eslicarbazepine acetate (300 μM) on Na^+^ current characteristics in MDA-MB-231 and HEK-Na_v_1.5 cells.^1^

| **A. MDA-MB-231 cells** |  |  |  |  |
| --- | --- | --- | --- | --- |
| *Parameter* | *Control* | *ESL* | *P value* | *N* |
| V<sub>thres</sub> (mV) | -45.7 ± 1.7 | -45.0 ± 1.4 | 0.58 | 13 |
| V<sub>peak</sub> (mV) | 3.1 ± 2.1 | -3.9 ± 2.7 | 0.056 | 13 |
| Activation V<sub>1/2</sub> (mV) | -19.3 ± 1.4 | -22.0 ± 1.5 | 0.095 | 12 |
| Activation k (mV) | 10.6 ± 0.7 | 9.3 ± 0.8 | 0.076 | 12 |
| Inactivation V<sub>1/2</sub> (mV) | -80.6 ± 0.7 | -86.7 ± 1.2 | <0.001 | 13 |
| Inactivation k (mV) | -4.8 ± 0.4 | -7.4 ± 1.7 | 0.139 | 13 |
| Peak current density at -10 mV (pA/pF) | -14.8 ± 3.9 | -8.0 ± 2.5 | <0.001 | 13 |
| Persistent current density at -10 mV (pA/pF) | -0.15 ± 0.05 | -0.02 ± 0.07 | 0.13 | 12 |
| T<sub>p</sub> at -10 mV (ms) | 2.1 ± 0.2 | 1.9 ± 0.2 | <0.01 | 13 |
| $\tau_f$ at -10 mV (ms) | 1.3 ± 0.1 | 1.3 ± 0.2 | 0.954 | 13 |
| $\tau_s$ at -10 mV (ms) | 10.0 ± 2.3 | 6.9 ± 2.0 | 0.289 | 13 |
| $\tau_r$ (ms) | 6.0 ± 0.5 | 8.7 ± 0.7 | <0.05 | 10 |
| **B. HEK-Na<sub>v</sub>1.5 cells** |  |  |  |  |
| *Parameter* | *Control* | *ESL* | *P value* | *N* |
| V<sub>thres</sub> (mV) | -55.0 ± 1.7 | -54.0 ± 2.2 | 0.758 | 10 |
| V<sub>peak</sub> (mV) | -26.0 ± 2.2 | -24.0 ± 4.3 | 0.591 | 10 |
| Activation V<sub>1/2</sub> (mV) | -39.4 ± 1.3 | -44.2 ± 1.8 | <0.05 | 10 |
| Activation k (mV) | 5.3 ± 1.3 | 3.8 ± 0.7 | 0.361 | 10 |
| Inactivation V<sub>1/2</sub> (mV) | -78.2 ± 2.5 | -88.3 ± 2.7 | <0.001 | 10 |
| Inactivation k (mV) | -6.9 ± 0.4 | -9.8 ± 0.7 | <0.001 | 10 |
| Peak current density at -10 mV (pA/pF) | -154.4 ± 24.0 | -33.1 ± 4.7 | <0.001 | 12 |
| Persistent current density at -10 mV (pA/pF) | -0.61 ± 0.15 | -0.12 ± 0.05 | <0.01 | 12 |
| T<sub>p</sub> at -10 mV (ms) | 1.4 ± 0.2 | 1.9 ± 0.2 | <0.001 | 14 |
| $\tau_f$ at -10 mV (ms) | 0.9 ± 0.1 | 1.2 ± 0.1 | <0.001 | 12 |
| $\tau_s$ at -10 mV (ms) | 6.6 ± 0.8 | 20.8 ± 8.5 | 0.128 | 12 |
| $\tau_r$ (ms) | 4.5 ± 0.4 | 7.1 ± 0.6 | <0.001 | 10 |

<sup>1</sup>ESL: eslicarbazepine acetate (300 μM); V<sub>thres</sub>: threshold voltage for activation; V<sub>peak</sub>: voltage at which current was maximal; V<sub>1/2</sub>: half (in)activation voltage; k: slope factor for (in)activation; T<sub>p</sub>: time to peak current; $\tau_f$ : fast time constant of inactivation; $\tau_s$ : slow time constant of inactivation; $\tau_r$ : time constant of recovery from inactivation. The holding potential was -120 mV. Results are mean ± SEM. Statistical comparisons were made with paired t-tests.

**Table 2.** Effect of S-licarbazepine (300 μM) on Na^+^ current characteristics in MDA-MB-231 and HEK-Na_v_1.5 cells.^1^

| **A. MDA-MB-231 cells** |  |  |  |  |
| --- | --- | --- | --- | --- |
| *Parameter* | *Control* | *S-Lic* | *P value* | *N* |
| V<sub>thres</sub> (mV) | -34.4 ± 2.0 | -35.7 ± 2.0 | 0.603 | 7 |
| V<sub>peak</sub> (mV) | 11.43 ± 4.4 | 10.0 ± 4.9 | 0.818 | 7 |
| Activation V<sub>1/2</sub> (mV) | -12.9 ± 1.3 | -13.7 ± 1.4 | 0.371 | 7 |
| Activation k (mV) | 11.0 ± 0.5 | 11.9 ± 0.8 | 0.520 | 7 |
| Inactivation V<sub>1/2</sub> (mV) | -71.8 ± 2.5 | -76.8 ± 2.2 | <0.05 | 7 |
| Inactivation k (mV) | -6.8 ± 0.9 | -6.0 ± 1.2 | 0.302 | 7 |
| Peak current density at -10 mV (pA/pF) | -12.0 ± 3.1 | -6.9 ± 2.5 | <0.001 | 9 |
| Persistent current density at -10 mV (pA/pF) | -1.3 ± 0.4 | -0.6 ± 0.2 | <0.05 | 9 |
| T<sub>p</sub> at -10 mV (ms) | 4.5 ± 0.4 | 5.1 ± 0.7 | 0.103 | 9 |
| τ<sub>f</sub> at -10 mV (ms) | 3.8 ± 1.1 | 3.2 ± 0.4 | 0.553 | 7 |
| τ<sub>s</sub> at -10 mV (ms) | 25.7 ± 7.0 | 27.1 ± 12.0 | 0.920 | 7 |
| τ<sub>r</sub> (ms) | 6.8 ± 0.4 | 13.5 ± 1.0 | <0.01 | 7 |
| **B. HEK-Na<sub>v</sub>1.5 cells** |  |  |  |  |
| *Parameter* | *Control* | *S-Lic* | *P value* | *N* |
| V<sub>thres</sub> (mV) | -50.0 ± 1.9 | -51.3 ± 3.5 | 0.598 | 9 |
| V<sub>peak</sub> (mV) | -18.0 ± 4.2 | -30.0 ± 5.6 | <0.001 | 9 |
| Activation V<sub>1/2</sub> (mV) | -32.8 ± 3.1 | -40.5 ± 3.4 | <0.01 | 9 |
| Activation k (mV) | 5.9 ± 0.9 | 4.5 ± 1.1 | <0.05 | 9 |
| Inactivation V<sub>1/2</sub> (mV) | -75.9 ± 2.6 | -79.3 ± 4.1 | 0.116 | 9 |
| Inactivation k (mV) | -6.5 ± 0.4 | -8.1 ± 0.5 | <0.05 | 9 |
| Peak current density at -10 mV (pA/pF) | -140.9 ± 26.8 | -77.2 ± 17.0 | <0.001 | 13 |
| Persistent current density at -10 mV (pA/pF) | -0.9 ± 0.2 | -0.5 ± 0.2 | <0.05 | 11 |
| T<sub>p</sub> at -10 mV (ms) | 1.8 ± 0.5 | 2.3 ± 0.6 | <0.01 | 13 |
| τ<sub>f</sub> at -10 mV (ms) | 1.0 ± 0.04 | 1.3 ± 0.06 | <0.001 | 11 |
| τ<sub>s</sub> at -10 mV (ms) | 6.3 ± 0.5 | 7.3 ± 0.5 | <0.05 | 11 |
| τ<sub>r</sub> (ms) | 5.7 ± 0.7 | 8.0 ± 1.2 | <0.01 | 10 |

<sup>1</sup>S-Lic: S-licarbazepine (300 µM); V<sub>thres</sub>: threshold voltage for activation; V<sub>peak</sub>: voltage at which current was maximal; V<sub>1/2</sub>: half (in)activation voltage; k: slope factor for (in)activation; T<sub>p</sub>: time to peak current; τ<sub>f</sub>: fast time constant of inactivation; τ<sub>s</sub>: slow time constant of inactivation; τ<sub>r</sub>: time constant of recovery from inactivation. The holding potential was -120 mV. Results are mean ± SEM. Statistical comparisons were made with paired t-tests.

### 3.2 Effect of eslicarbazepine acetate and S-licarbazepine on voltage dependence of activation and inactivation

We next investigated the effect of ESL (300 µM) and S-Lic (300 µM) on the I-V relationship in MDA-MB-231 and HEK-Na_v_1.5 cells. A V_h_ of −120 mV was used for subsequent analyses to ensure that the elicited currents were sufficiently large for analysis of kinetics and voltage dependence, particularly for MDA-MB-231 cells, which display smaller peak Na^+^ currents (Tables 1, 2). Neither ESL nor S-Lic had any effect on the threshold voltage for activation (Figure 4A-D; Tables 1, 2). ESL also had no effect on the voltage at current peak in either cell line (Figure 4A-D; Tables 1, 2).

**Figure 4.**
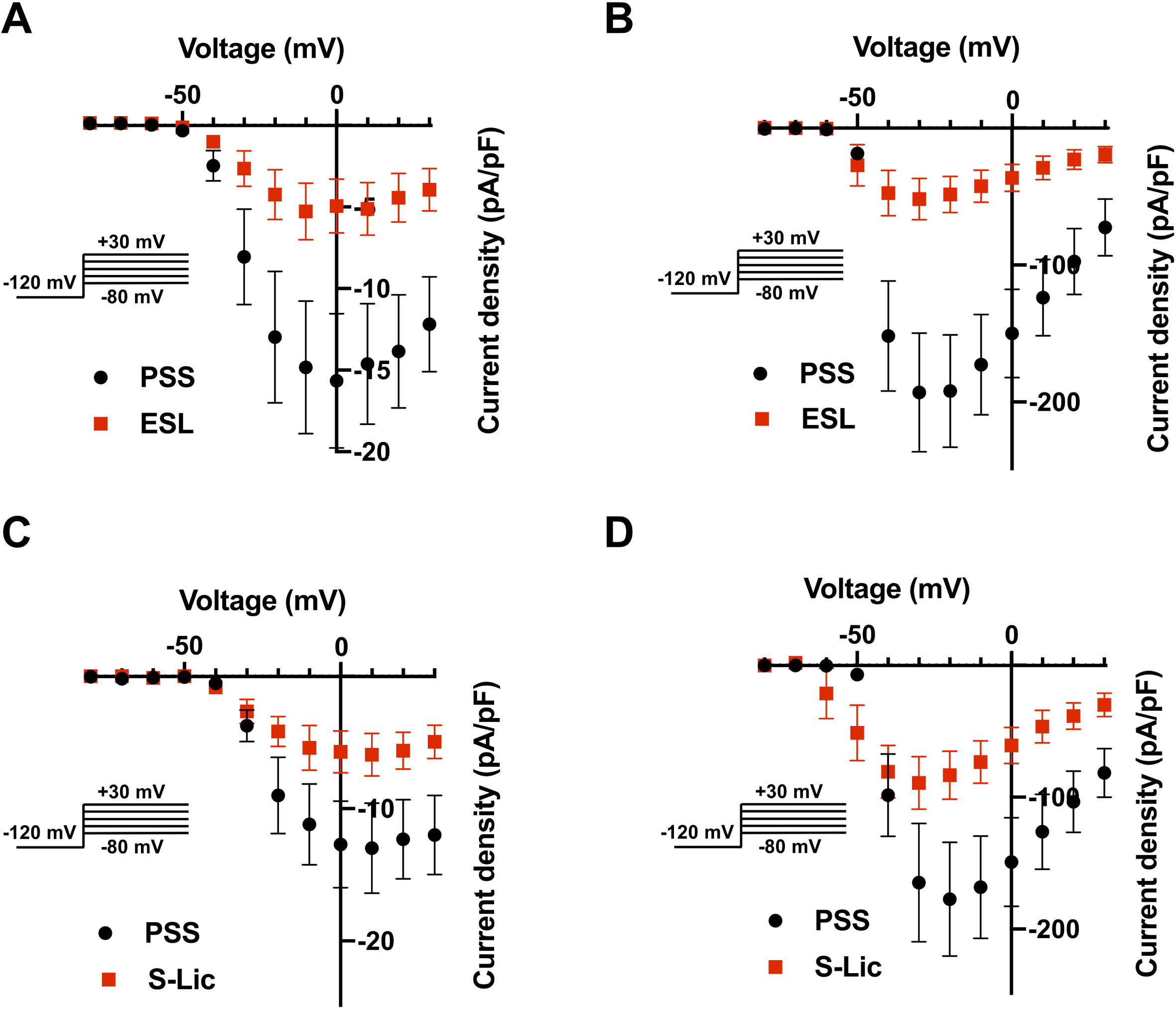
Effect of eslicarbazepine acetate and S-licarbazepine on the current-voltage relationship. (A) Current-voltage (I-V) plots of Na^+^ currents in MDA-MB-231 cells in physiological saline solution (PSS; black circles) and in eslicarbazepine acetate (ESL; 300 μM; red squares). (B) (I-V) plots of Na^+^ currents in HEK-Na_v_1.5 cells in PSS (black circles) and ESL (300 μM; red squares). (C) I-V plots of Na^+^ currents in MDA-MB-231 cells in PSS (black circles) and S-licarbazepine (S-Lic; 300 μM; red squares). (D) I-V plots of Na^+^ currents in HEK-Na_v_1.5 cells in PSS (black circles) and S-Lic (300 μM; red squares). Currents were elicited using 10 mV depolarising steps from −80 to +30 mV for 30 ms, from a holding potential of −120 mV. Results are mean ± SEM (n = 7-13).

Although S-Lic had no effect on voltage at current peak in MDA-MB-231 cells, it was significantly hyperpolarised in HEK-Na_v_1.5 cells from −18.0 ± 4.2 mV to −30.0 ± 5.6 mV (P < 0.001; n = 9; paired t test; Figure 4A-D; Tables 1, 2). ESL had no significant effect on the half-activation voltage (V½) or slope factor (k) for activation in MDA-MB-231 cells (Figure 5A; Table 1). The activation k in HEK-Na_v_1.5 cells was also unchanged but the activation V½ was significantly hyperpolarised by ESL from −39.4 ± 1.3 to −44.2 ± 1.8 mV (P < 0.05; n = 10; paired t test; Figure 5B; Table 1). S-Lic also had no significant effect on the activation V½ or k in MDA-MB-231 cells (Figure 5C; Table 2). However, the V½ of activation in HEK-Na_v_1.5 cells was significantly hyperpolarised from −32.8 ± 3.1 mV to −40.5 ± 3.4 mV (P < 0.01; n = 9; paired t test; Figure 5D; Table 2) and k changed from 5.9 ± 0.9 mV to 4.5 ± 1.1 mV (P < 0.05; n = 9; paired t test; Figure 5D; Table 2).

**Figure 5.**
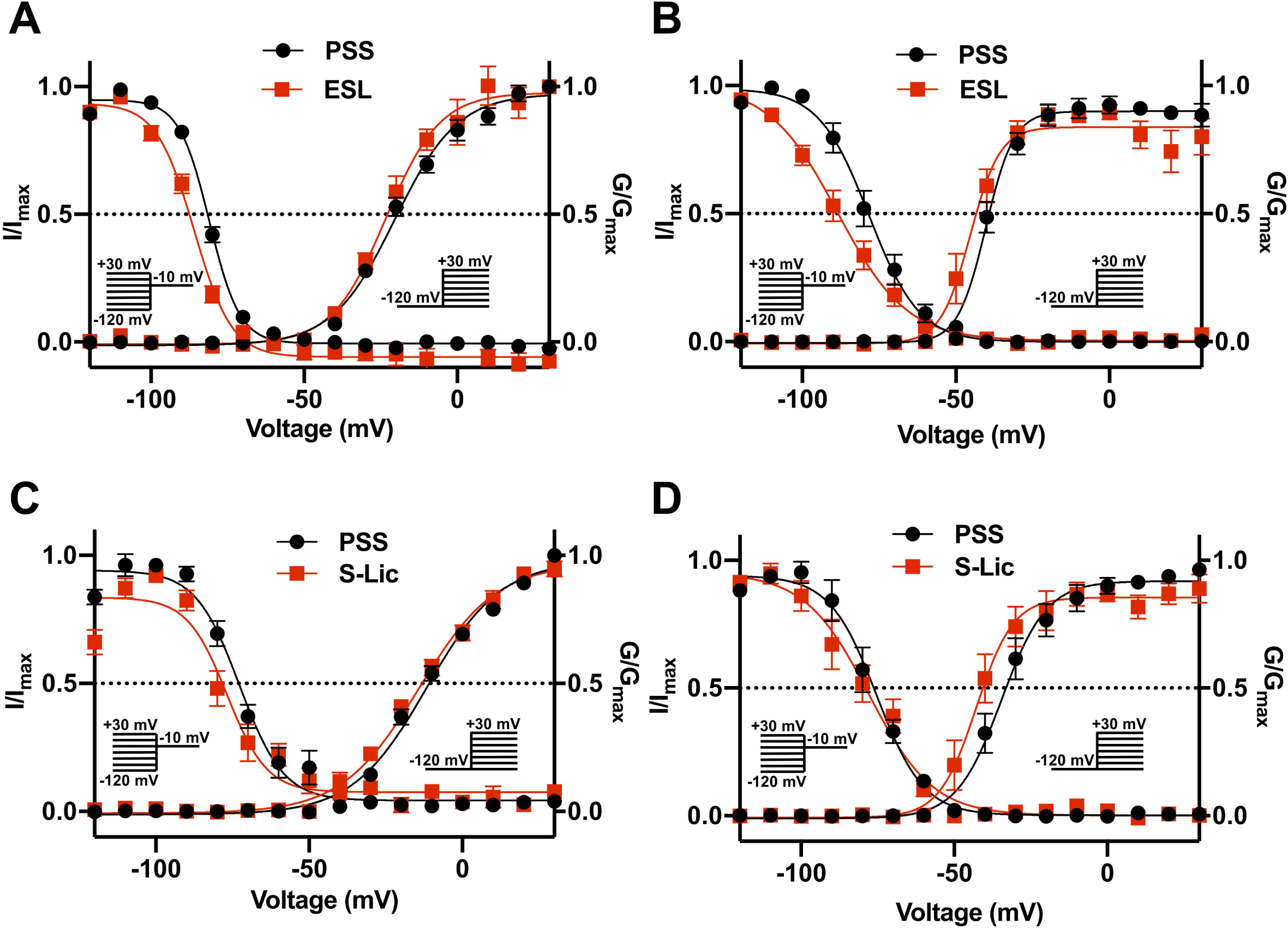
Effect of eslicarbazepine acetate and S-licarbazepine on activation and steady-state inactivation. (A) Activation and steady-state inactivation in MDA-MB-231 cells in physiological saline solution (PSS; black circles) and in eslicarbazepine acetate (ESL; 300 μM; red squares). (B) Activation and steady-state inactivation in HEK-Na_v_1.5 cells in PSS (black circles) and ESL (300 μM; red squares). (C) Activation and steady-state inactivation in MDA-MB-231 cells in PSS (black circles) and S-licarbazepine (S-Lic; 300 μM; red squares). (D) Activation and steady-state inactivation in HEK-Na_v_1.5 cells in PSS (black circles) and S-Lic (300 μM; red squares). For activation, normalised conductance (G/G_max_) was calculated from the current data and plotted as a function of voltage. For steady-state inactivation, normalised current (I/I_max_), elicited by 50 ms test pulses at −10 mV following 250 ms conditioning voltage pulses between −120 mV and +30 mV, applied from a holding potential of −120 mV, was plotted as a function of the prepulse voltage. Results are mean ± SEM (n = 7-13). Activation and inactivation curves are fitted with Boltzmann functions.

As regards steady-state inactivation, in MDA-MB-231 cells, ESL significantly hyperpolarised the inactivation V½ from −80.6 ± 0.7 mV to −86.7 ± 1.2 mV (P < 0.001; n = 13; paired t test) without affecting inactivation k (Figure 5A; Table 1). ESL also hyperpolarised the inactivation V½ in HEK-Na_v_1.5 cells from −78.2 ± 2.5 mV to −88.3 ± 2.7 mV (P < 0.001; n = 10; paired t test), and changed the inactivation k from −6.9 ± 0.4 mV to −9.8 ± 0.7 mV (P < 0.001; n = 10; paired t test; Figure 5B; Table 1). S-Lic also significantly hyperpolarised the inactivation V½ in MDA-MB-231 cells from −71.8 ± 2.5 mV to −76.8 ± 2.2 mV (P < 0.05; n = 7; paired t test) without affecting inactivation k (Figure 5C; Table 2). However, the inactivation V½ in HEK-Na_v_1.5 cells was not significantly altered by S-Lic, although the inactivation k significantly changed from −6.5 ± 0.4 mV to −8.1 ± 0.5 mV (P < 0.05; n = 9; paired t test; Figure 5D; Table 2). In summary, both ESL and S-Lic affected various aspects of the voltage dependence characteristics of Na_v_1.5 in MDA-MB-231 and HEK-Na_v_1.5 cells, predominantly hyperpolarising the voltage dependence of inactivation.

### 3.3 Effect of eslicarbazepine acetate and S-licarbazepine on activation and inactivation kinetics

We next studied the effect of both compounds on kinetics of activation and inactivation. In MDA-MB-231 cells, ESL (300 μM) significantly accelerated the time to peak current (T_p_), upon depolarisation from −120 mV to −10 mV, from 2.1 ± 0.2 ms to 1.9 ± 0.2 ms (P < 0.01; n = 13; paired t test; Table 1). However, in HEK-Na_v_1.5 cells, ESL significantly slowed T_p_ from 1.4 ± 0.2 ms to 1.5 ± 0.2 ms (P < 0.001; n = 14; paired t test; Table 1). S-Lic (300 μM) had no significant effect on T_p_ in MDA-MB-231 cells but significantly slowed T_p_ in HEK-Na_v_1.5 cells from 1.8 ± 0.5 ms to 2.3 ± 0.6 ms (P < 0.01; n = 13; paired t test; Table 2).

To study effects on inactivation kinetics, the current decay following depolarisation from −120 mV to −10 mV was fitted to a double exponential function to derive fast and slow time constants of inactivation (τ_f_ and τ_s_). Neither ESL nor S-Lic had any significant effect on τ_f_ or τ_s_ in MDA-MB-231 cells (Tables 1, 2). However, in HEK-Na_v_1.5 cells, ESL significantly slowed τ_f_ from 0.9 ± 0.1 ms to 1.2 ± 0.1 ms (P < 0.001; n = 12; paired t test; Table 1) and slowed τ_s_ from 6.6 ± 0.8 ms to 20.8 ± 8.5 ms, although this was not statistically significant. S-Lic significantly slowed τ_f_ from 1.0 ± 0.04 ms to 1.3 ± 0.06 ms (P < 0.001; n = 11; paired t test; Table 2) and τ_s_ from 6.3 ± 0.5 ms to 7.3 ± 0.5 ms (P < 0.05; n = 11; paired t test; Table 2). In summary, both ESL and S-Lic elicited various effects on kinetics in MDA-MB-231 and HEK-Na_v_1.5 cells, predominantly slowing activation and inactivation.

### 3.4 Effect of eslicarbazepine acetate and S-licarbazepine on recovery from fast inactivation

To investigate the effect of ESL and S-Lic on channel recovery from fast inactivation, we subjected cells to two depolarisations from V_h_ of −120 mV to 0 mV, changing the interval between these in which the channels were held at −120 mV to facilitate recovery. Significance was determined by fitting a single exponential curve to the normalised current/time relationship and calculating the time constant (τ_r_). In MDA-MB-231 cells, ESL (300 μM) significantly slowed τ_r_ from 6.0 ± 0.5 ms to 8.7 ± 0.7 ms (P < 0.05; n = 10; paired t test; Figure 6A, Table 1). Similarly, in HEK-Na_v_1.5 cells, ESL significantly slowed τ_r_ from 4.5 ± 0.4 ms to 7.1 ± 0.6 ms (P < 0.001; n = 10; paired t test; Figure 6B, Table 1). S-Lic (300 μM) also significantly slowed τ_r_ in MDA-MB-231 cells from 6.8 ± 0.4 ms to 13.5 ± 1.0 ms (P < 0.01; n = 7; paired t test; Figure 6C, Table 2). Finally, S-Lic also significantly slowed τ_r_ in HEK-Na_v_1.5 cells from 5.7 ± 0.7 ms to 8.0 ± 1.2 ms (P < 0.01; n = 10; paired t test; Figure 6D, Table 2). In summary, both ESL and S-Lic slowed recovery from fast inactivation of Na_v_1.5.

**Figure 6.**
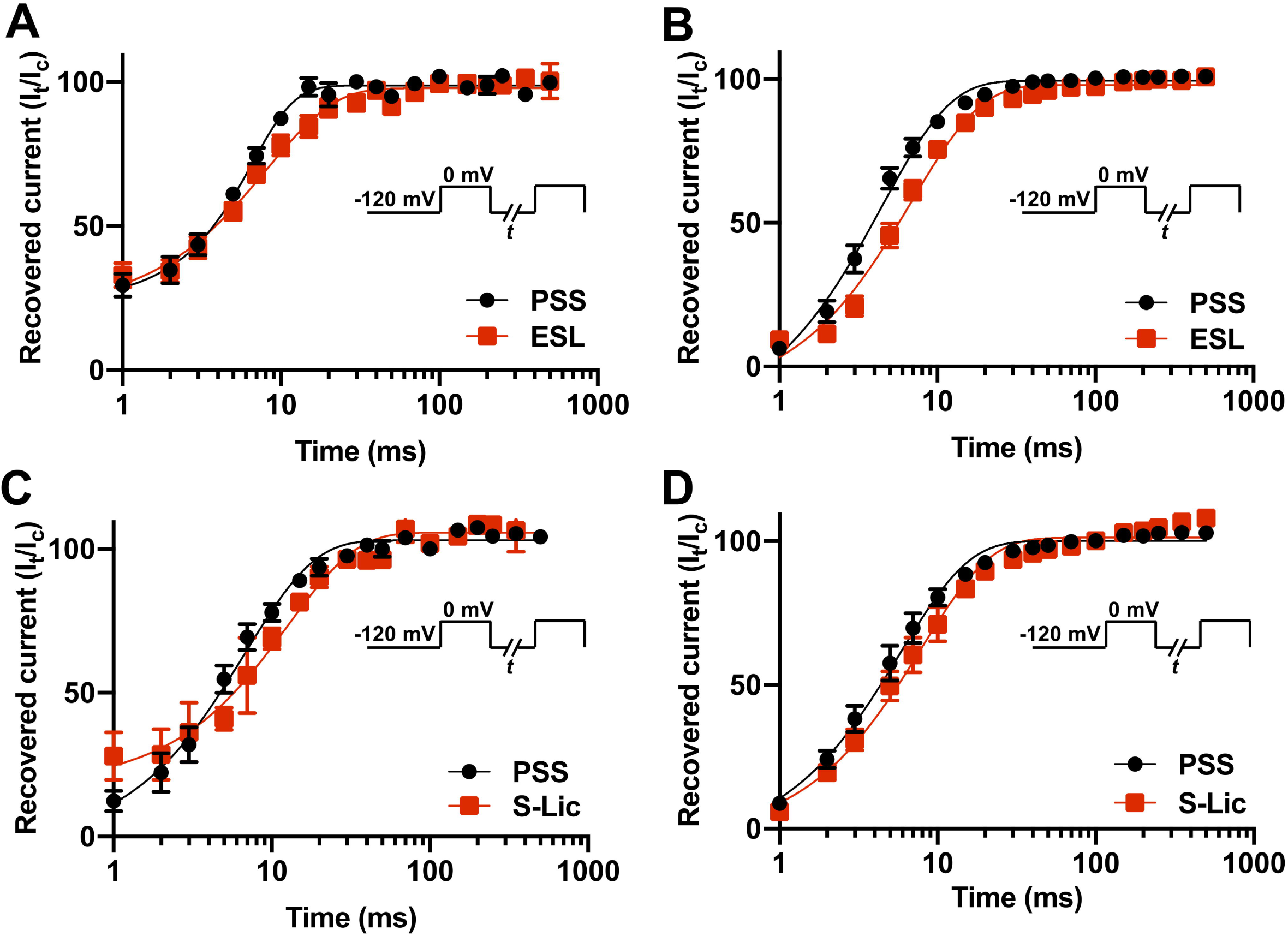
Effect of eslicarbazepine acetate and S-licarbazepine on recovery from inactivation. (A) Recovery from inactivation in MDA-MB-231 cells in physiological saline solution (PSS; black circles) and in eslicarbazepine acetate (ESL; 300 μM; red squares). (B) Recovery from inactivation in HEK-Na_v_1.5 cells in PSS (black circles) and ESL (300 μM; red squares). (C) Recovery from inactivation in MDA-MB-231 cells in PSS (black circles) and S-licarbazepine (S-Lic; 300 μM; red squares). (D) Recovery from inactivation in HEK-Na_v_1.5 cells in PSS (black circles) and S-Lic (300 μM; red squares). The fraction recovered (I_t_/I_c_) was determined by a 25 ms pulse to 0 mV (I_c_), followed by a recovery pulse to −120 mV for 1-500 ms, and a subsequent 25 ms test pulse to 0 mV (I_t_), applied from a holding potential of −120 mV, and plotted as a function of the recovery interval. Data are fitted with single exponential functions which are statistically different between control and drug treatments in all cases. Results are mean ± SEM (n = 7-10).

## 4 Discussion

In this study, we have shown that ESL and its active metabolite S-Lic inhibit the transient and persistent components of Na^+^ current carried by Na_v_1.5. We show broadly similar effects in MDA-MB-231 cells, which express endogenous Na_v_1.5 (29, 30, 45), and in HEK-293 cells over-expressing Na_v_1.5. Notably, both compounds were more effective when V_h_ was set to −80 mV than at −120 mV, suggestive of depolarised state-dependent binding. In addition, the inhibitory effect of ESL was reversible whereas inhibition by S-Lic was less so. As regards voltage-dependence, both ESL and S-Lic shifted activation and steady-state inactivation curves, to varying extents in the two cell lines, in the direction of more negative voltages. ESL and S-Lic had various effects on activation and inactivation kinetics, generally slowing the rate of inactivation. Finally, recovery from fast inactivation of Na_v_1.5 was significantly slowed by both ESL and S-Lic.

To our knowledge, this is the first time that the effects of ESL and S-Lic have specifically been tested on the Na_v_1.5 isoform. A strength of this study is that both the prodrug (ESL) and the active metabolite (S-Lic) were tested using two independent cell lines, one endogenously expressing Na_v_1.5, the other stably over-expressing Na_v_1.5. MDA-MB-231 cells also express Na_v_1.7, although this isoform is estimated to be responsible for only ∼9 % of the total VGSC current (30, 45). MDA-MB-231 cells also express endogenous β1, β2 and β4 subunits (47-49). MDA-MB-231 cells predominantly express the developmentally regulated ‘neonatal’ Na_v_1.5 splice variant, which differs from the ‘adult’ variant over-expressed in the HEK-Na_v_1.5 cells by seven amino acids located in the extracellular linker between transmembrane segments 3 and 4 of domain 1 (30, 42, 45). Notably, however, there were no consistent differences in effect of either ESL or S-Lic between the MDA-MB-231 and HEK-Na_v_1.5 cells, suggesting that the neonatal vs. adult splicing event, and/or expression of endogenous β subunits, does not impact on sensitivity of Na_v_1.5 to these compounds. This finding contrasts another report showing different sensitivity of the neonatal and adult Na_v_1.5 splice variants to the amide local anaesthetics lidocaine and levobupivacaine (44). Our findings suggest that the inhibitory effect of S-Lic on Na_v_1.5 is less reversible than that of ESL. This may be explained by the differing chemical structures of the two molecules possibly enabling S-Lic to bind the target with higher affinity than ESL. Most VGSC-targeting anticonvulsants, including phenytoin, lamotrigine and carbamazepine, block the pore by binding via aromatic-aromatic interaction to a tyrosine and phenylalanine located in the S6 helix of domain 4 (50). However, S-Lic has been proposed to bind to a different site given that it was found to block the pore predominantly during slow inactivation (10). Alternatively, the hydroxyl group present on S-Lic (but not ESL) may become deprotonated, potentially trapping it in the cytoplasm.

The findings presented here broadly agree with *in vitro* concentrations used elsewhere to study effects of ESL and S-Lic on Na^+^ currents. For example, using a V_h_ of −80 mV, 300 µM ESL was shown to inhibit peak Na^+^ current by 50 % in N1E-115 neuroblastoma cells expressing Na_v_1.1, Na_v_1.2, Na_v_1.3, Na_v_1.6 and Na_v_1.7 (20). S-Lic (250 µM) also blocks peak Na^+^ current by ∼50 % in the same cell line (10). In addition, S-Lic (300 µM) reduces persistent Na^+^ current by ∼25 % in acutely isolated murine hippocampal CA1 neurons expressing Na_v_1.1, Na_v_1.2 and Na_v_1.6 (21-24). Similar to the present study, ESL was shown to hyperpolarise the voltage-dependence of steady-state inactivation in N1E-115 cells (20). On the other hand, similar to our finding in HEK-Na_v_1.5 cells, S-Lic has no effect on steady-state inactivation in N1E-115 cells (10). Again, in agreement with our own findings for Na_v_1.5, S-Lic slows recovery from inactivation in N1E-115 cells (10). These observations suggest that the sensitivity of Na_v_1.5 to ESL and S-Lic is broadly similar to that reported for neuronal VGSCs. In support of this, Na_v_1.5 shares the same conserved residues proposed for Na_v_1.2 to interact with ESL (Figure 7) (51).

**Figure 7.**
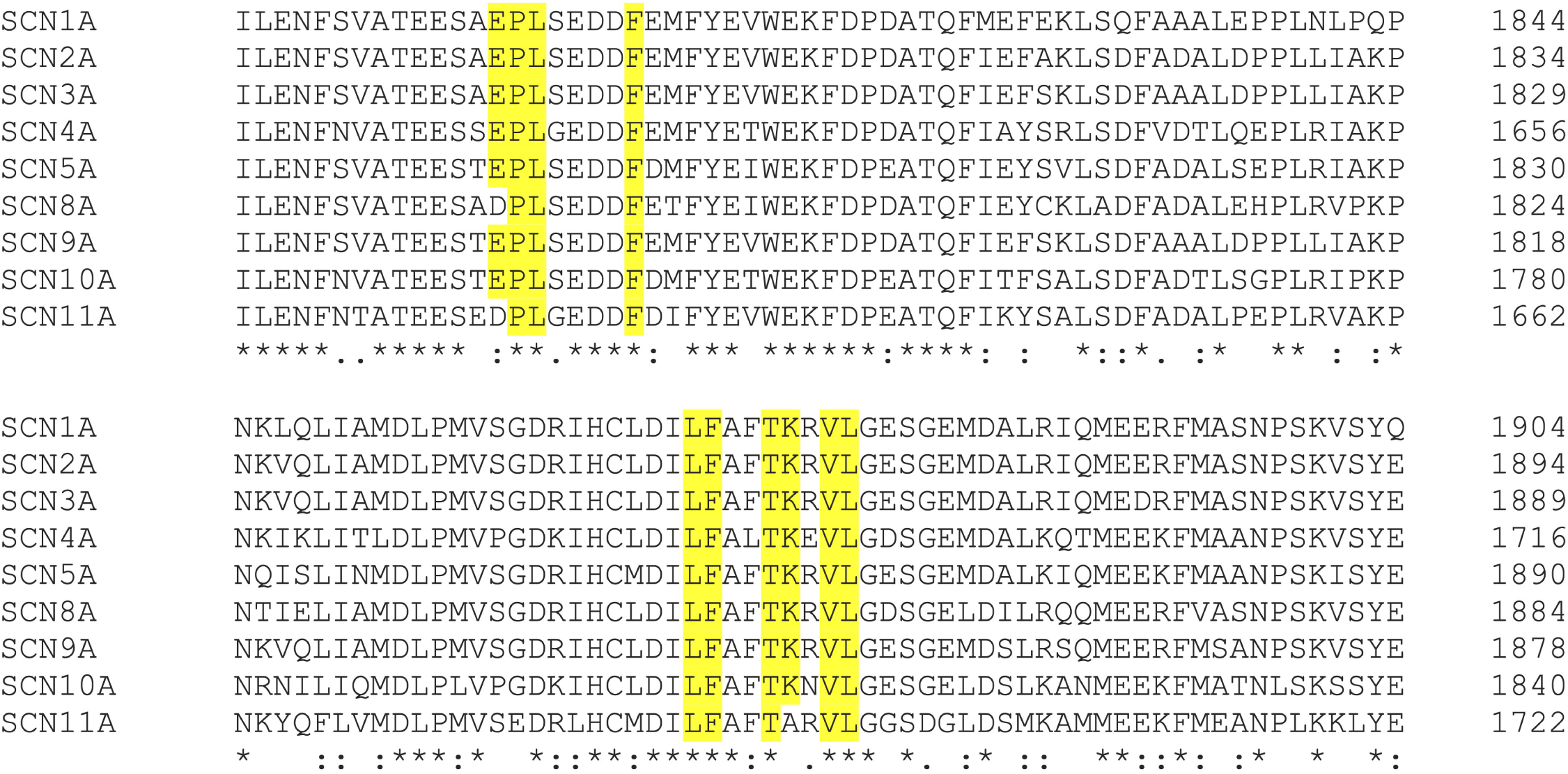
Clustal alignment of amino acid sequences of Na_v_1.1-Na_v_1.9 (*SCN1A-SCN11A*). ESL was proposed previously (51) to interact with the highlighted amino acids in Na_v_1.2. An alignment of Na_v_1.2 (UniProtKB - Q99250 (SCN2A_HUMAN)) with Na_v_1.1 (UniProtKB - P35498 (SCN1A_HUMAN)), Na_v_1.3 (UniProtKB - Q9NY46 (SCN3A_HUMAN)), Na_v_1.4 (UniProtKB - P35499 (SCN4A_HUMAN)), Na_v_1.5 (UniProtKB - Q14524 (SCN5A_HUMAN)) Na_v_1.6 (UniProtKB - Q9UQD0 (SCN8A_HUMAN)), Na_v_1.7 (UniProtKB - Q15858 (SCN9A_HUMAN)), Na_v_1.8 (UniProtKB - Q9Y5Y9 (SCN10A_HUMAN)), and Na_v_1.9 (UniProtKB - Q9UI33 (SCN11A_HUMAN)) shows that the interacting amino acids highlighted in yellow are conserved between Na_v_1.2 and Na_v_1.5, along with most other isoforms. Asterisks indicate conserved residues. Colon indicates conservation between groups of strongly similar properties - scoring > 0.5 in the Gonnet PAM 250 matrix. Period indicates conservation between groups of weakly similar properties - scoring ≤ 0.5 in the Gonnet PAM 250 matrix.

Notably, the concentrations used in this study are at or above those achieved in clinical use (e.g. ESL 1200 mg once daily gives a peak plasma concentration of ∼100 µM) (10). However, it has been argued that the relatively high concentrations which have been previously tested *in vitro* are clinically relevant given that S-Lic has a high (50:1) lipid:water partition co-efficient and thus would be expected to reside predominantly in the tissue membrane fraction *in vivo* (15). Our study suggests that a clinically relevant plasma concentration (100 µM) would inhibit peak and persistent Na_v_1.5 currents. Future work investigating the dose-dependent effects of ESL and S-Lic would be useful to aid clinical judgements.

The data presented here raise several implications for clinicians. The observed inhibition of Na_v_1.5 is worthy of note when considering cardiac function in patients receiving ESL (13). Although the QT interval remains unchanged for individuals on ESL treatment, prolongation of the PR interval has been observed (27). Further work is required to establish whether the basis for this PR prolongation is indeed via Na_v_1.5 inhibition. In addition, it would be of interest to investigate the efficacy of ESL and S-Lic in the context of heritable arrhythmogenic mutations in *SCN5A*, as well as the possible involvement of the β subunits (24, 26, 52, 53). The findings presented here are also relevant in the context of Na_v_1.5 expression in carcinoma cells (54). Given that cancer cells have a relatively depolarised V_m_, it is likely that Na_v_1.5 is mainly in the inactivated state with the persistent Na^+^ current being functionally predominant (55, 56). Increasing evidence suggests that persistent Na^+^ current carried by Na_v_1.5 in cancer cells contributes to invasion and several studies have shown that other VGSC inhibitors reduce metastasis in preclinical models (29-35, 57). Thus, use-dependent inhibition by ESL would ensure that channels in malignant cells are particularly targeted, raising the possibility that it could be used as an anti-metastatic agent (43). This study therefore paves the way for future investigations into ESL and S-Lic as potential invasion inhibitors.

## 5 Author Contributions

TL, SC and WB contributed to the conception and design of the work. TL, LB and WB contributed to acquisition, analysis, and interpretation of data for the work. TL, SC and WB contributed to drafting the work and revising it critically for important intellectual content. All authors approved the final version of the manuscript.

## 6 Abbreviations

ESL: eslicarbazepine acetate;
HEK-Na_v_1.5: HEK-293 cells stably expressing Na_v_1.5;
I-V: current-voltage;
k: slope factor;
PSS: physiological saline solution;
S-Lic: S-licarbazepine
T_p_: time to peak current;
τ_f_: fast time constant of inactivation;
τ_s_: slow time constant of inactivation;
τ_r_: time constant of recovery from inactivation;
VGSC: voltage-gated Na^+^ channel;
V_m_: membrane potential;
V_h_: holding potential;
V_peak_: voltage at which current was maximal;
V_rev_: reversal potential;
V_thres_: threshold voltage for activation;
V_1/2_: half-activation voltage.

## 7 Acknowledgements

This work was supported by Cancer Research UK (A25922) and Breast Cancer Now (2015NovPhD572).

## 8 Conflict of interest statement

The authors declare that the research was conducted in the absence of any commercial or financial relationships that could be construed as a potential conflict of interest.

## 9 Data availability statement

The datasets used and/or analysed during the current study are available from the corresponding author on reasonable request.

## Notes

### Competing Interest Statement

The authors have declared no competing interest.

