## Supplementary figure legends for "Inhibitory effect of eslicarbazepine acetate and S-licarbazepine on Na_v_1.5 channels"

**Supplementary Figure 1.** Effect of 0.45% DMSO on VGSC current-voltage relationship and gating in MDA-MB-231 cells. (A) Current-voltage (I-V) plots of Na^+^ currents in MDA-MB-231 cells in physiological saline solution (PSS; black circles) and in PSS with 0.45% DMSO (0.45% DMSO; green squares). Currents were elicited using 10 mV depolarising steps from -80 to +30 mV for 30 ms, from a holding potential of -120 mV. Results are mean ± SEM (n = 13-17). (B) Activation and steady-state inactivation in physiological saline solution (PSS; black circles) and in PSS with 0.45% DMSO (0.45% DMSO; green squares). For activation, normalised conductance (G/G_max_) was calculated from the current data and plotted as a function of voltage. For steady-state inactivation, normalised current (I/I_max_), elicited by 50 ms test pulses at -10 mV following 250 ms conditioning voltage pulses between -120 mV and +30 mV, applied from a holding potential of -120 mV, was plotted as a function of the prepulse voltage. Results are mean ± SEM (n = 10-13). Activation and inactivation curves are fitted with Boltzmann functions.
