## Supplementary Table 1 for "Inhibitory effect of eslicarbazepine acetate and S-licarbazepine on Na_v_1.5 channels"

**Supplementary Table 1A.** Effect of eslicarbazepine acetate (100 μM) on peak and persistent Na^+^ current in MDA-MB-231 and HEK-Na_v_1.5 cells.

| **A. MDA-MB-231 cells** |  |  |  |  |
| --- | --- | --- | --- | --- |
| *Parameter* | *Control* | *ESL* | *P value* | *N* |
| Peak current density at -10 mV, V_h_ -120 mV (pA/pF) | -22.1 ± 13.5 | -11.6 ± 7.9 | <0.05 | 7 |
| Peak current density at -10 mV, V_h_ -80 mV (pA/pF) | -7.1 ± 4.1 | -2.1 ± 2.0 | < 0.05 | 7 |
| Persistent current density at -10 mV, V_h_ -120 mV (pA/pF) | -0.5 ± 0.3 | -0.4 ± 0.2 | 0.277 | 7 |
| **B. HEK-Na_v_1.5 cells** |  |  |  |  |
| *Parameter* | *Control* | *ESL* | *P value* | *N* |
| Peak current density at -10 mV, V_h_ -120 mV (pA/pF) | -158.4 ± 85.7 | -77.7 ± 51.3 | <0.01 | 8 |
| Peak current density at -10 mV, V_h_ -80 mV (pA/pF) | -59.0 ± 50.7 | -12.2 ± 11.9 | <0.05 | 8 |
| Persistent current density at -10 mV, V_h_ -120 mV (pA/pF) | -1.0 ± 0.3 | -0.4 ± 0.1 | <0.001 | 8 |

^1^ESL: eslicarbazepine acetate (100 µM). Results are mean ± SEM. Statistical comparisons were made with paired t-tests.

**Supplementary Table 1B.** Effect of S-licarbazepine (100 μM) on peak and persistent Na^+^ current in MDA-MB-231 and HEK-Na_v_1.5 cells.

| **A. MDA-MB-231 cells** |  |  |  |  |
| --- | --- | --- | --- | --- |
| *Parameter* | *Control* | *S-Lic* | *P value* | *N* |
| Peak current density at -10 mV, V_h_ -120 mV (pA/pF) | -17.2 ± 8.7 | -12.3 ± 7.4 | 0.084 | 8 |
| Peak current density at -10 mV, V_h_ -80 mV (pA/pF) | -7.8 ± 4.7 | -3.5 ± 2.6 | <0.05 | 8 |
| Persistent current density at -10 mV, V_h_ -120 mV (pA/pF) | -0.6 ± 0.3 | -0.4 ± 0.2 | <0.01 | 8 |
| **B. HEK-Na_v_1.5 cells** |  |  |  |  |
| *Parameter* | *Control* | *S-Lic* | *P value* | *N* |
| Peak current density at -10 mV, V_h_ -120 mV (pA/pF) | -108.5 ± 20.3 | - 75.6 ± 30.9 | <0.05 | 8 |
| Peak current density at -10 mV, V_h_ -80 mV (pA/pF) | -30.2 ± 0.9 | -11.8 ± 1.3 | <0.001 | 8 |
| Persistent current density at -10 mV, V_h_ -120 mV (pA/pF) | -0.5 ± 0.1 | -0.3 ± 0.1 | <0.05 | 7 |

^1^S-Lic: S-licarbazepine (100 µM). Results are mean ± SEM. Statistical comparisons were made with paired t-tests.
