## Supplementary figures and images for "Inhibitory effect of eslicarbazepine acetate and S-licarbazepine on Na_v_1.5 channels"

### Supplementary figure 1

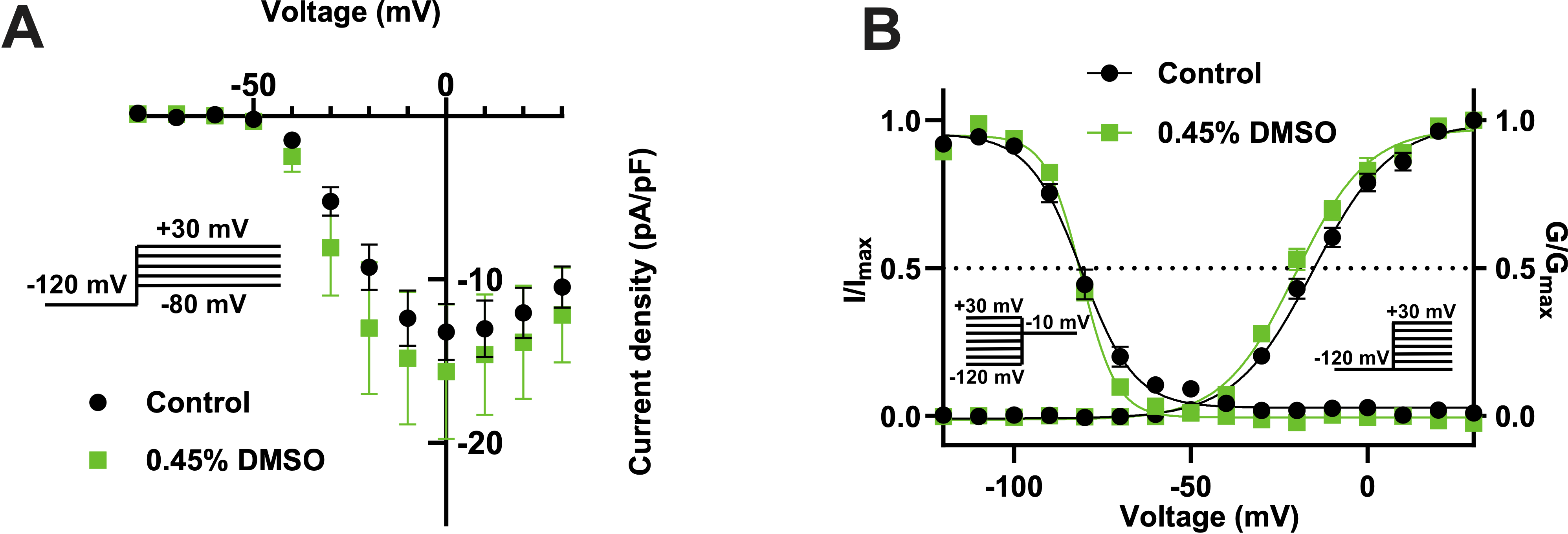

### Supplementary figure 2

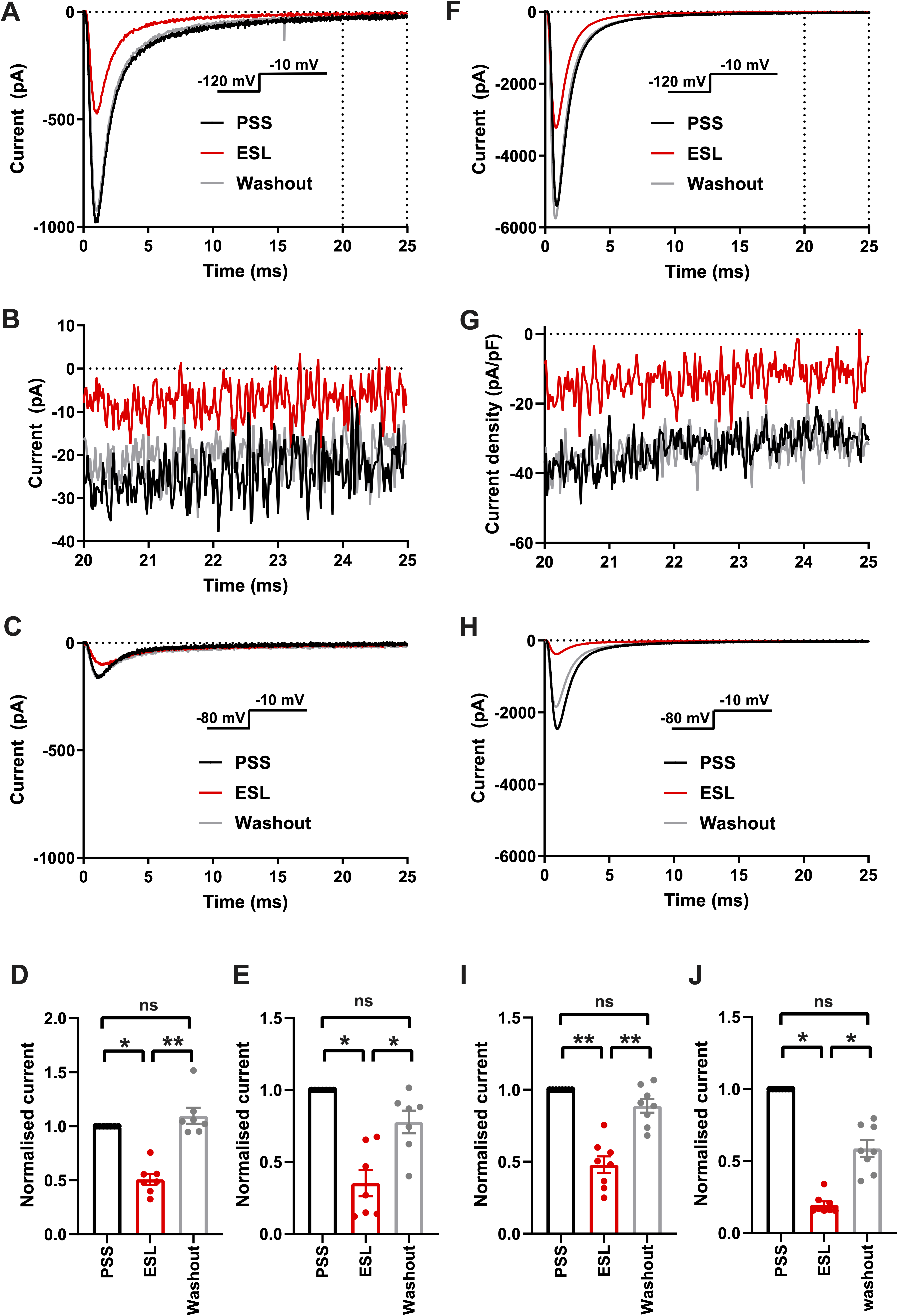

### Supplementary figure 3

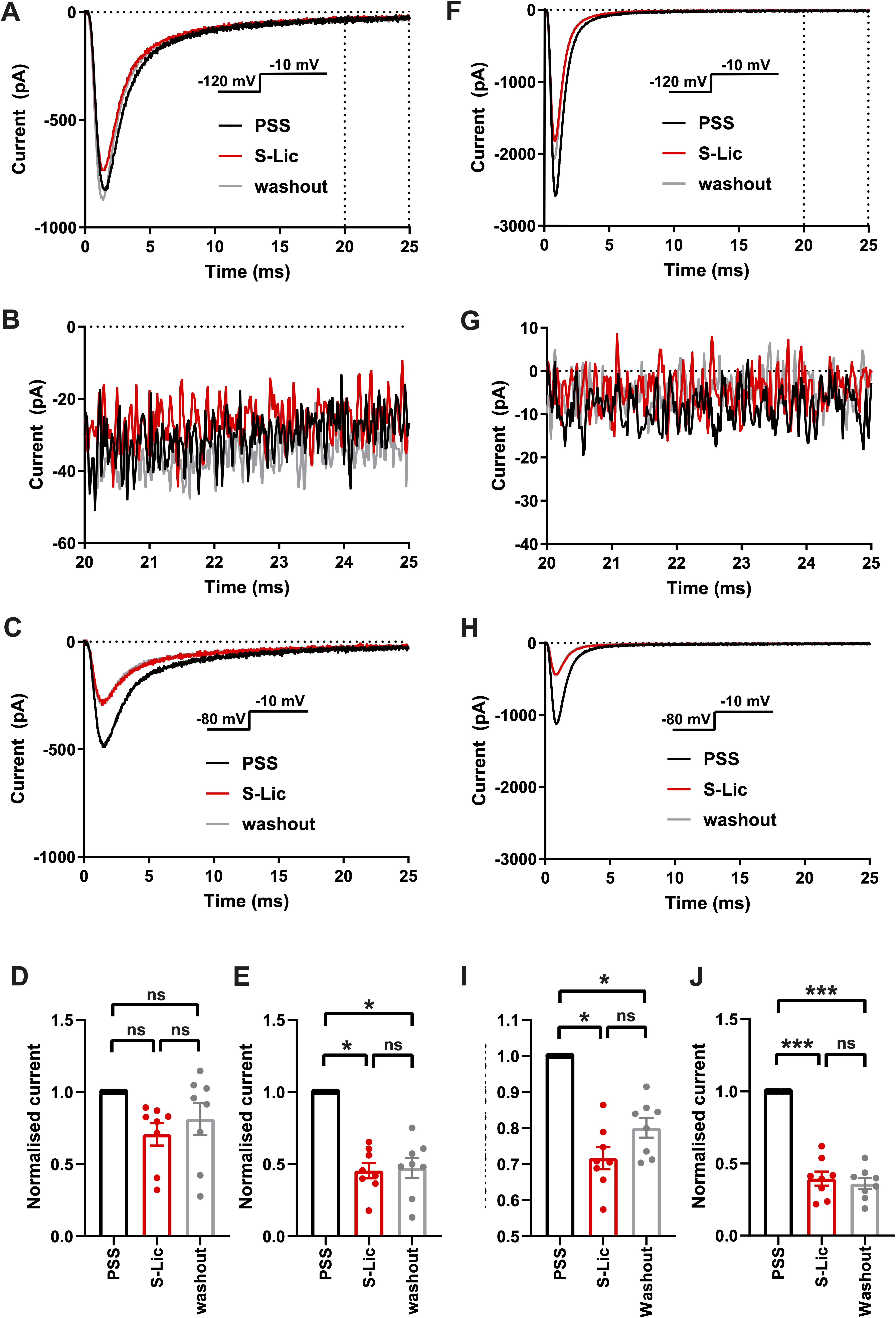
